# SNAREs and Synaptotagmin cooperatively determine the Ca^2+^ sensitivity of neurotransmitter release in fixed stoichiometry modules

**DOI:** 10.1101/2021.06.22.449527

**Authors:** Zachary A. McDargh, Ben O’Shaughnessy

## Abstract

Neurotransmitter release is accomplished by a multi-component machinery including the membrane-fusing SNARE proteins and Ca2+-sensing Synaptotagmin molecules. However, the Ca^2+^ sensitivity of release was found to increase or decrease with more or fewer SNARE complexes at the release site, respectively, while the cooperativity is unaffected (Acuna et al., 2014; Arancillo et al., 2013), suggesting that there is no simple division of labor between these two components. To examine the mechanisms underlying these findings, we developed molecular dynamics simulations of the neurotransmitter release machinery, with variable numbers of Synaptotagmin molecules and assembled SNARE complexes at the release site. Ca^2+^ uncaging simulations showed that increasing the number of SNARE complexes at fixed stoichiometric ratio of Synaptotagmin to SNAREs increased the Ca^2+^ sensitivity without affecting the cooperativity. The physiological cooperativity of ~4-5 was reproduced with 2-3 Synaptotagmin molecules per SNARE complex, suggesting that Synaptotagmin and SNAREs cooperate in fixed stoichiometry modules. In simulations of action potential-evoked release, increased numbers of Synaptotagmin-SNARE modules increased release probability, consistent with experiment. Our simulations suggest that the final membrane fusion step is driven by SNARE complex-mediated entropic forces, and by vesicle-tethering forces mediated by the long Synaptotagmin linker domains. In consequence, release rates are increased when more SNARE complexes and Synaptotagmin monomers are present at the fusion site.

## Introduction

Neurotransmitter (NT) release at a chemical synapse is accomplished by a many-component protein machinery that holds synaptic vesicles in a primed state in preparation for release, senses Ca^2+^ influx through voltage-gated Ca^2+^ channels, and fuses the membranes in response, releasing vesicle contents through the resultant fusion pore. Synchronization of NT release with stimulus is essential for many cognitive functions (1) and relies on a molecular machinery that must achieve two functions: first, it inhibits membrane fusion in basal Ca^2+^ concentrations (termed “clamping”) and releases this inhibition in elevated Ca^2+^ concentrations (2–5). Second, once the release machinery is unclamped, it fuses synaptic vesicle and plasma membranes on sub-millisecond time scales (6).

The clamping and Ca^2+^-triggered unclamping functions are thought to be achieved by Synaptotagmin 1 (Syt) (3, 4, 7, 8) (although Complexin has also been proposed as a clamp (9, 10)). Syt comprises two Ca^2+^-binding C2 domains, connected to a vesicle-associated trans-membrane domain (TMD) by a 60-residue linker domain (LD). Deletion of Syt leads to a dramatic increase in spontaneous release frequency in mice and *Drosophila*, suggesting that Syt clamps release at basal Ca^2+^ concentrations (2–4, 8, 11, 12). Syt deletion also causes release to shift from a synchronous to an asynchronous mode (8, 13, 14), suggesting that Syt activates fusion in response to elevated Ca^2+^ concentrations. The mechanisms of these functions are not well-understood. Syt may achieve clamping by inhibiting SNARE zippering in basal Ca^2+^ concentrations (12, 15). Syt has been observed to assemble into ring-like oligomers *in vitro* that spontaneously disassemble in physiological Ca^2+^ concentrations (16, 17) which could clamp release by sterically separating the vesicle and plasma membranes (18, 19). Syt mutations selectively inhibiting oligomerization abolished Ca^2+^-sensitive clamping both *in vitro* and *in vivo* (18–21).

The second function of the NT release machinery, membrane fusion, is accomplished by the neuronal SNARE proteins VAMP, SNAP-25, and Syntaxin (22, 23); together, these proteins assemble from N-terminus to C-terminus into a four-helix bundle called a SNAREpin. Many details of the mechanism of SNARE-mediated fusion remain unclear. The energy released by SNAREpin assembly is often thought to be used to overcome the ~25 – 35 *kT* energetic barrier to fusion (24). According to this picture, the minimum number of SNAREs required for fusion is determined by a comparison of the zippering energy with the activation barrier to fusion. However, no machinery that is capable of transducing the SNARE assembly energy into membrane energy has been established. EPR and NMR spectroscopy showed that the SNARE LDs are unstructured and flexible (25–27), suggesting they are unable to store the zippering energy as bending energy. An alternative model is that fusion is driven by entropic forces originating from SNARE-SNARE and SNARE-membrane interactions that press the vesicle and plasma membranes together (28, 29). According to this model, any number of SNAREs can achieve fusion, but the SNARE-mediated fusion rate increases exponentially with additional SNARE complexes at the fusion site.

However, several studies have found that it is not possible to neatly divide the NT release machinery into these two non-overlapping parts, as genetic and pharmacological alterations of the fusion machinery have led to correlated effects on the Ca^2+^ sensitivity of NT release, illustrating a coupling between the Ca^2+^-sensing and membrane-fusing components of the NT release machinery. Acuna et al. (30) and Gerber et al. (31) found that NT release is enhanced at mouse calyx of Held synapses and cortical neurons, respectively, by a constitutively open mutant of the t-SNARE Syntaxin, which increased the number of assembled SNARE complexes at the fusion site. The vesicle release probability, NT release rate and, surprisingly, apparent Ca^2+^ sensitivity of release were all increased, while the apparent Ca^2+^ cooperativity of release, i.e. the scaling of the vesicle release rate as a function of [Ca^2+^], was unaffected. Conversely, Arancillo et al. found that mutations reducing Syntaxin 1 expression in presynaptic nerve terminals lowered the vesicle release probability and the Ca^2+^ sensitivity of release (32). Calyx of Held presynaptic terminals treated with botulism neurotoxin A, botulism neurotoxin C1, or tetanus neurotoxin, which cleave SNAP-25, Syntaxin, and VAMP, respectively, also displayed lower Ca^2+^ sensitivity (33), with no change in cooperativity. These results indicate that the apparent Ca^2+^ sensitivity of release is partially determined by elements of the release machinery more typically associated with fusion.

Here, we performed coarse-grained (CG) molecular dynamics (MD) simulations of NT release, varying the number of Syt monomers and SNARE complexes at the fusion site in order to investigate the mechanism of this coupling. In simulations, we assume that Syt molecules are initially assembled into a ring-like oligomer that clamps release prior to arrival of Ca^2+^, and that SNARE complexes bound to this ring via the primary interface identified in the Syt-SNARE complex crystal structure (34). Our simulations reproduce the dramatic enhancement of NT release with additional SNARE complexes observed experimentally (30–32). In simulations of Ca^2+^ uncaging, we found that increasing the number of SNARE complexes at the fusion site led to an increase in EPSC amplitude and the Ca^2+^ sensitivity of release. In simulations, the experimentally observed conservation of the Ca^2+^ cooperativity of release with additional SNAREs required the stoichiometry of the release machinery to be fixed, as simulations predicted that the cooperativity is proportional to the stoichiometric ratio of Syt molecules to SNARE complexes at the release site. The experimental cooperativity of ~4 was reproduced with 2-3 Syt molecules per SNARE complex, suggesting that *in vivo* Syt molecules and SNARE complexes form modules with 2-3 Syt molecules per SNARE complex. In simulations of AP-evoked release, we found a dramatic increase in release probability with additional Syt-SNARE modules, consistent with experiment. This enhancement of NT release is driven by entropic forces mediating membrane fusion together with tethering forces mediated by the Syt LD that pull the vesicle and PM together. These forces increase with more SNARE complexes and more Syt molecules, respectively (28, 29, 35).

## Results

### Model of the NT release machinery

In order to investigate the Ca^2+^-sensitive clamping and membrane-fusing functions of the NT release machinery, as well as the coupling between them, we developed a minimal model of the release machinery, incorporating the core elements of machinery for both tasks. We therefore present CG molecular dynamics simulations of 40 nm synaptic vesicles hosting *N*_Syt_ copies of Syt 1, docked to a planar plasma membrane (PM) by a collection of *N*_SNARE_ SNARE complexes, Fig. 1. SNARE complexes in our simulations zipper and unzipper dynamically.

**Figure 1:**
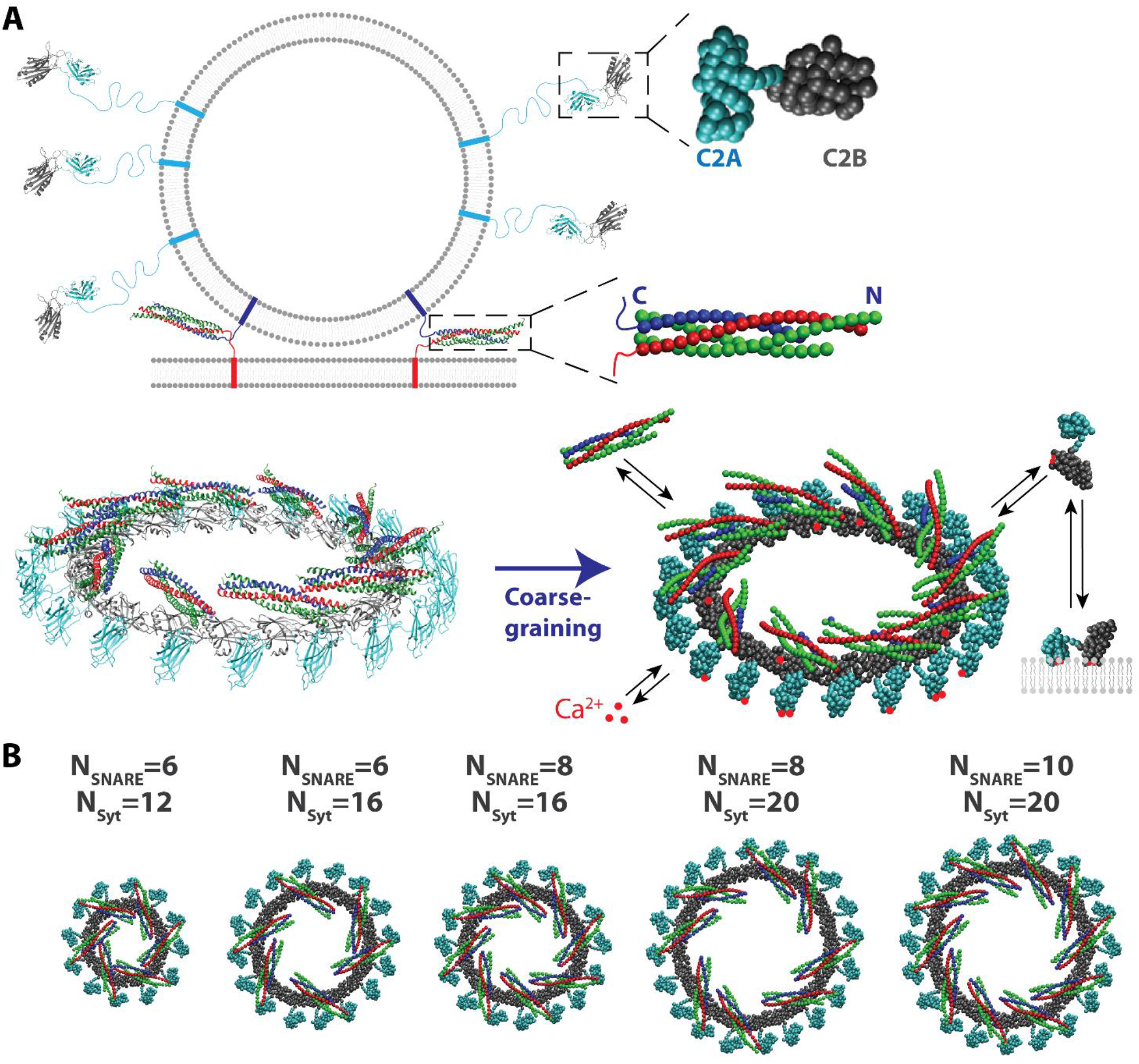
Coarse-grained molecular dynamics simulations. **(A)** Top: Coarse-grained (CG) representations of Synaptotagmin 1 (Syt) and the neuronal SNARE complex, with VAMP shown in blue, Syntaxin in red, and SNAP-25 in green. Bottom: atomistic representation (left) and coarse-grained representation (right) of the Syt-SNARE ring. The structure of the Syt ring was reconstructed from cryo-electron micrographs of helical Syt polymers (16). SNARE complexes were docked to the ring via the primary interface located on the Syt C2B domain identified in ref. (34). Syt-SNARE rings are interposed between the vesicle and PM in the initial condition of our simulations. Syt-SNARE and Syt-membrane bonds dynamically form and dissociate throughout the simulations. **(B)** Our simulations vary the number of SNARE complexes and Syt molecules at the release site, with *N*_SNARE_ ranging from 6 to 10, and *N*_Syt_ ranging from 12 to 20, with the requirement that *N*_Syt_ ≥ 2*N*_SNARE_ in order to avoid steric clashes between SNARE complexes in the ring.

The Syt molecules are initially arranged in a ring-shaped oligomer (16, 17, 36, 37) to which SNARE complexes are bound via the primary interface identified in crystal structures of the Syt-SNARE complex (34). *In vitro* studies have found that Syt spontaneously assembles into oligomers on negatively charged lipid membranes, which are disassembled in physiological Ca^2+^ concentrations, leading to the hypothesis that Syt rings might clamp fusion before the arrival of an action potential (AP) at an axon terminal (16, 17, 36).

We simulated both the situation before arrival of an AP, when presynaptic intracellular [Ca^2+^] is fixed at 0.1 μM (38, 39), and the situation when intracellular [Ca^2+^] is elevated, triggering membrane fusion and NT release. To determine the system’s response to both steady-state and transiently elevated [Ca^2+^], we considered elevation of [Ca^2+^] from Ca^2+^ uncaging and AP-evoked influx of Ca^2+^ through voltage gated Ca^2+^ channels.

Our simulations incorporate dynamic assembly and disassembly of the SNARE complexes. Furthermore, we incorporate dynamic binding and dissociation of Syt-SNARE, Syt-Syt, and Syt-PM bonds, as well as binding of Ca^2+^ ions to the Ca^2+^-binding loops of the Syt C2A and C2B domains, Fig. 1.

#### CG Models

Our simulations employed highly coarse-grained models of the vesicle and plasma membranes, Syt 1, and the neuronal SNARE complex (28, 29, 35). See Supporting Information for model parameters.

The vesicle and plasma membrane are represented by non-deformable, uniformly charged sphere and plane, respectively, with charge density determined by the known lipid composition of synaptic vesicles and the presynaptic PM, Table S4. Membrane fusion is assumed to occur when the interaction energy between the membranes exceeds the activation barrier to fusion (24), here assumed to be *E*_fusion_ = 20 *kT*, slightly lower than the 25 – 35 *kT* measured experimentally for pure phosphatidylcholine (PC) membranes, an assumption made to account for the highly fusogenic composition of the vesicle and plasma membranes, including high concentrations of cholesterol and phosphatidylethanolamine (PE), both of which promote membrane fusion (40, 41).

Our CG protein models were generated by mapping groups of residues from crystal structures of the SNARE complex (PDB ID 3hd7 (42)) and Syt 1 (PDB ID 5ccg (34)) to CG beads. In alpha helix or beta strand regions of the protein, groups of four residues were mapped to a single bead, while in unstructured loops two residues were mapped to a bead. The structure of the globular domains of the proteins was fixed, so that the Syt C2AB domains and the assembled part of the SNARE complexes are non-deformable. The Syt 1 linker domain and unassembled parts of Syntaxin and VAMP are represented implicitly as worm-like chains using the free energy of worm-like chain extension (43).

#### SNARE assembly dynamics

The SNARE complex comprises 16 layers, numbered −7 to +8, which zipper from the N terminal end to the C terminal end as the complex assembles. Simulated disassembly events converted one bead of VAMP to four residues in the unstructured region connecting the SNARE motif to the trans-membrane domain (26). Disassembly events in the C-terminal domain of the SNARE complex (layers +5-+8) removed one bead from each helix of the SNARE complex (44).

Assembly and disassembly events occurred with rates 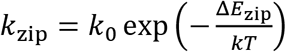 and *k*_unzip_= *k*_0_ = 10^6^ s^−1^, respectively, where the zippering energy Δ*E*_zip_ is the sum of contributions from the measured free energy landscape of SNARE assembly and the energetic cost of stretching the uncomplexed linker domain (44–46). See ref. (35) for details of the calculation of the zippering energy.

#### Reaction scheme

Our simulations incorporated a dynamic binding scheme representing the formation and dissociation of Syt-SNARE bonds via the primary interface (34, 47), Syt-Syt bonds (16, 19, 36), and Syt-plasma membrane bonds (48). For all of these interactions, a bond is formed when the separation between residues making up the binding interface on the two binding partners is less than a pre-defined capture distance, Table S1. Bonds then dissociate stochastically at a rate *k*_off_ specific to that interaction. The *k*_off_ values of the Syt-Syt and Syt-SNARE interactions are tuned to reproduce the experimentally measured dissociation constants (36, 47). The dissociation rate of Syt from the PM was taken from optical tweezers measurements (49). Following experimental observations that Syt oligomers are disassembled in physiological Ca^2+^ concentrations, we assume that Syt-Syt bonds dissociate upon Ca^2+^ binding to both C2B binding sites (16).

Lastly, we include the dynamic association and dissociation of Ca^2+^ ions with the Ca^2+^-binding loops of the Syt C2A and C2B domains. The intracellular [Ca^2+^] is assumed uniform in all simulations. Ca^2+^ ions bind to each Ca^2+^-binding site at a rate *k*_on_[Ca^2+^], where *k*_on_ = 2 × 10^8^ M^−1^s^−1^, and dissociate at a rate determined by the dissociation constant of the associated binding site, Table S1. The on rate *k*_on_ and the dissociation constant of the C2B Ca^2+^-binding sites are tuned to reproduce physiological characteristics of AP-evoked release at the calyx of Held: the ~0.2 ms delay time from the start of Ca^2+^-influx to the start of an EPSC, and the release probability per vesicle of ~0.1 (35, 50). See ref. (35) for details of the parameter tuning procedure.

### Simulations with only the SNARE components of the release machinery

Our model of the NT release machinery features two core components, Syt and the SNARE complex, respectively the core components of the Ca^2+^-sensitive clamping and membrane fusion machineries. We begin by examining the membrane-fusing SNARE protein machinery on its own in order to isolate its effects on release. We performed simulations of synaptic vesicles docked to planar plasma membranes omitting Syt, Fig. 1 & 2D, and measured the mean time required for fusion as a function of the number of SNARE complexes at the release site. These simulations are similar to those performed in our previous studies (28, 29), but assume a more realistic membrane composition for the vesicle membrane and the neuronal PM and use a more recent SNARE zippering landscape which showed that there is an energetic barrier to zippering the C-terminal end of the SNARE complex (44).

**Figure 2:**
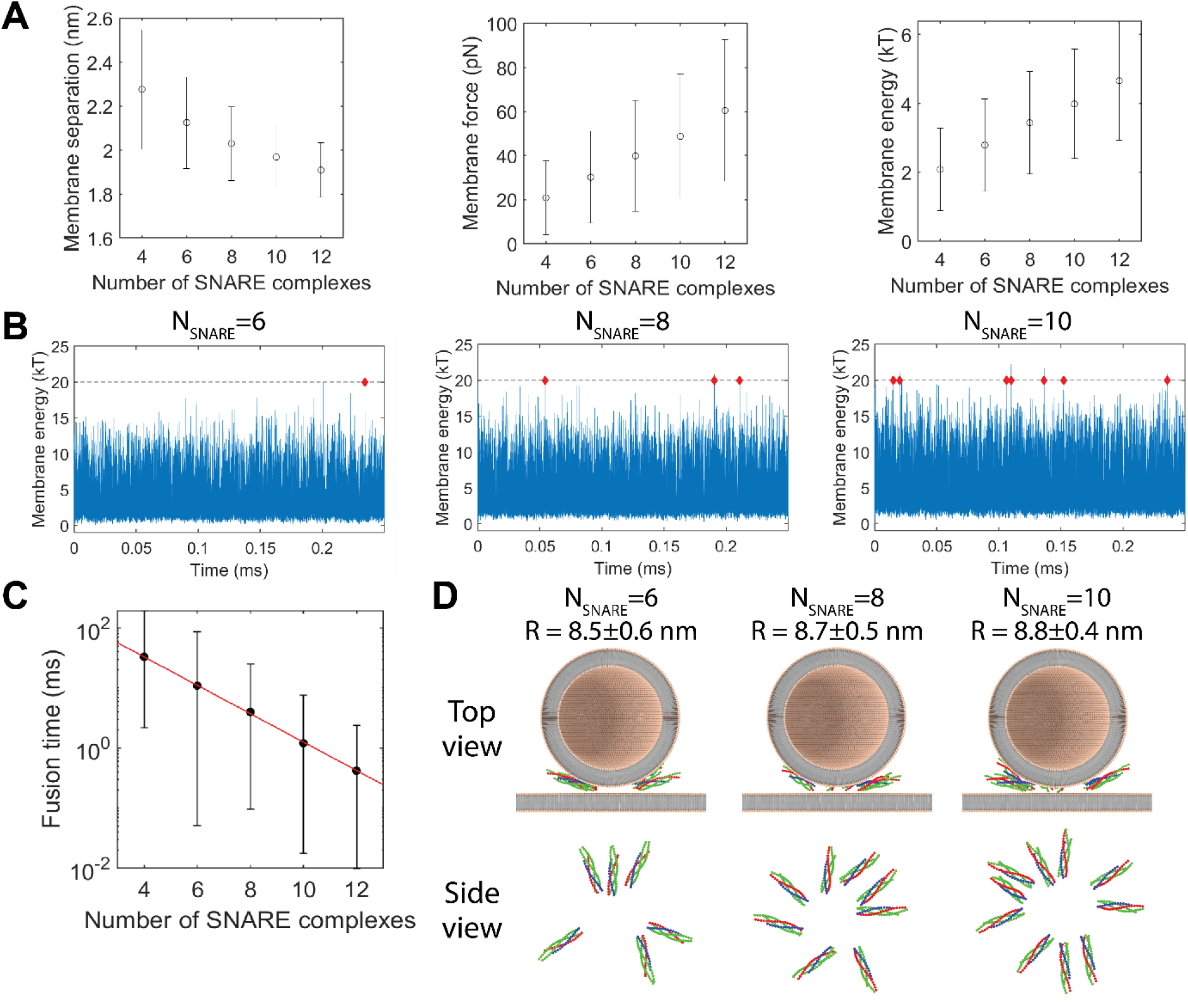
SNARE complexes self-assemble into rings that drive fusion through entropic forces. **(A)** Membrane separation (left), force (center), and energy (right) vs. number of SNARE complexes at the fusion site. Error bars are SD (taken from 10 simulations lasting 0.25 ms each). **(B)** Membrane energy vs. time, varying the number of SNARE complexes at the fusion site. Each plot shows the vesicle-PM interaction energy at every time step in the simulation. Dashed lines represent the activation barrier to fusion, taken to be 20 *kT* here. Red diamonds indicate times when the membrane energy exceeded the activation barrier, representing a fusion event. With more SNARE complexes, the membrane energy is increased, and has larger fluctuations, increasing the rate at which fusion events occur. **(C)** Fusion time vs. number of SNARE complexes. Black points represent the average time between fusion events observed in 10 simulations lasting 0.25 ms each; error bars are SD. Red line represents a best-fit exponential (*R*^2^ = 0.9999). **(D)** Simulation snapshots. With more SNARE complexes, SNARE rings are larger, pulling the vesicle closer to the PM due to the curvature of the vesicle. Membranes are illustrated using explicit lipids, while in simulations they were modeled as continuous surfaces.

It is commonly thought that the energy released by SNARE zippering is used directly to overcome the activation barrier to membrane fusion. We therefore first investigated the time required to zipper a collection of *N*_SNARE_ SNARE complexes. Our simulations show that SNARE complexes zipper independently, with no cooperativity between SNAREs.

The simulated SNARE complexes zippered rapidly, and the zippering time depended weakly on the number of SNARE complexes, Fig. S1. The time required for all SNARE motifs to become fully assembled to layer +8 increased from ~31 μs with 4 SNARE complexes to ~45 μs with 12 SNARE complexes, Fig. S1, much less than fusion time scales of ~1 ms (1). Our results are consistent with a mathematical model that we developed that assumes each SNARE complex zippers independently, and are inconsistent with a recent mathematical model that predicted the zippering time increases exponentially as a function of *N*_SNARE_ (51) (see Supporting Information section *Analytical model of SNARE zippering kinetics*).

As the SNARE complexes zippered, they spontaneously organized into a ring-like configuration, with their membrane proximal C terminal ends towards the center of the ring, and their membrane distal N terminal ends pointing radially outward, Fig. 2D, as observed in our previous studies (28, 29). This arrangement minimizes crowding by maximizing the freedom of the SNARE complexes to move laterally around the ring, thus maximizing entropy, similar to the occurrence of spontaneous nematic order in liquid crystals (52).

Entropy also favors a larger ring because, as the ring expands, the SNARE complexes are less crowded, and the SNARE complexes have greater freedom to rotate both in and out of the plane of the PM (28, 29). Thus, entropic forces arising from SNARE-SNARE and SNARE-membrane interactions drove the SNARE complexes outward, pulling the vesicle towards the plasma membrane. The radius of the ring and the vesicle-PM separation were set by the balance between entropic forces pushing the SNAREs outward and tension in the linker domains attaching the SNARE complexes to the vesicle and plasma membranes, pulling them inward.

The SNARE ring increased in radius with increasing numbers of SNARE complexes at the fusion site, Fig. 2D, showing that the entropic forces are stronger with more SNARE complexes. Due to the curvature of the vesicle membrane, radial expansion of the SNARE ring decreased the separation between the vesicle and the PM at the point of closest approach, thus increasing the interaction energy between the membranes and the associated force, Fig. 2A.

Fluctuations in the SNARE ring size drove large fluctuations in the membrane energy, eventually increasing the energy over the activation barrier to fusion, Fig. 2B. Increasing the number of SNARE complexes at the fusion site promoted ring expansion, accelerating fusion. The waiting time for fusion decreased exponentially as a function of the number of SNARE complexes at the fusion site *N*_SNARE_, Fig. 2B & C, consistent with our previous simulations (28, 29). With an activation barrier to fusion of 20 *kT*, fitting the observed fusion times to an exponential function yielded

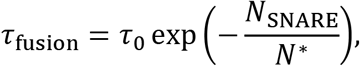

where *τ*_0_ = 290 ± 25 ms, and *N*\* = 1.84 ± 0.07. We note that this *N*\* value is slightly different from values we predicted previously (28, 29), likely due to the different membrane composition assumed here.

### Simulations of the full machinery: Ca^2+^-mediated unclamping and fusion

Prior to arrival of an AP at a synaptic terminal, release-competent vesicles are clamped. This clamp must remain stable at basal Ca^2+^ concentrations, and be removed in response to elevated Ca^2+^ concentrations. Thus we first asked if the Syt ring is able to serve as a Ca^2+^-sensitive fusion clamp. We simulated vesicles with 20 Syt monomers initially assembled into a ring, with 10 SNARE complexes docked to the Syt ring via the primary interface identified in the Syt-SNARE crystal structure (34), Fig. 3A. This number of Syt monomers is close to the average copy number per ring measured *in vitro* (36, 37) and the copy number hosted by synaptic vesicles (53, 54). [Ca^2+^] was held at either 0.1 μM, the basal value in cells, or 25 μM, comparable to the peak local [Ca^2+^] value reached in axon terminals during AP-evoked release (50, 55, 56).

**Figure 3:**
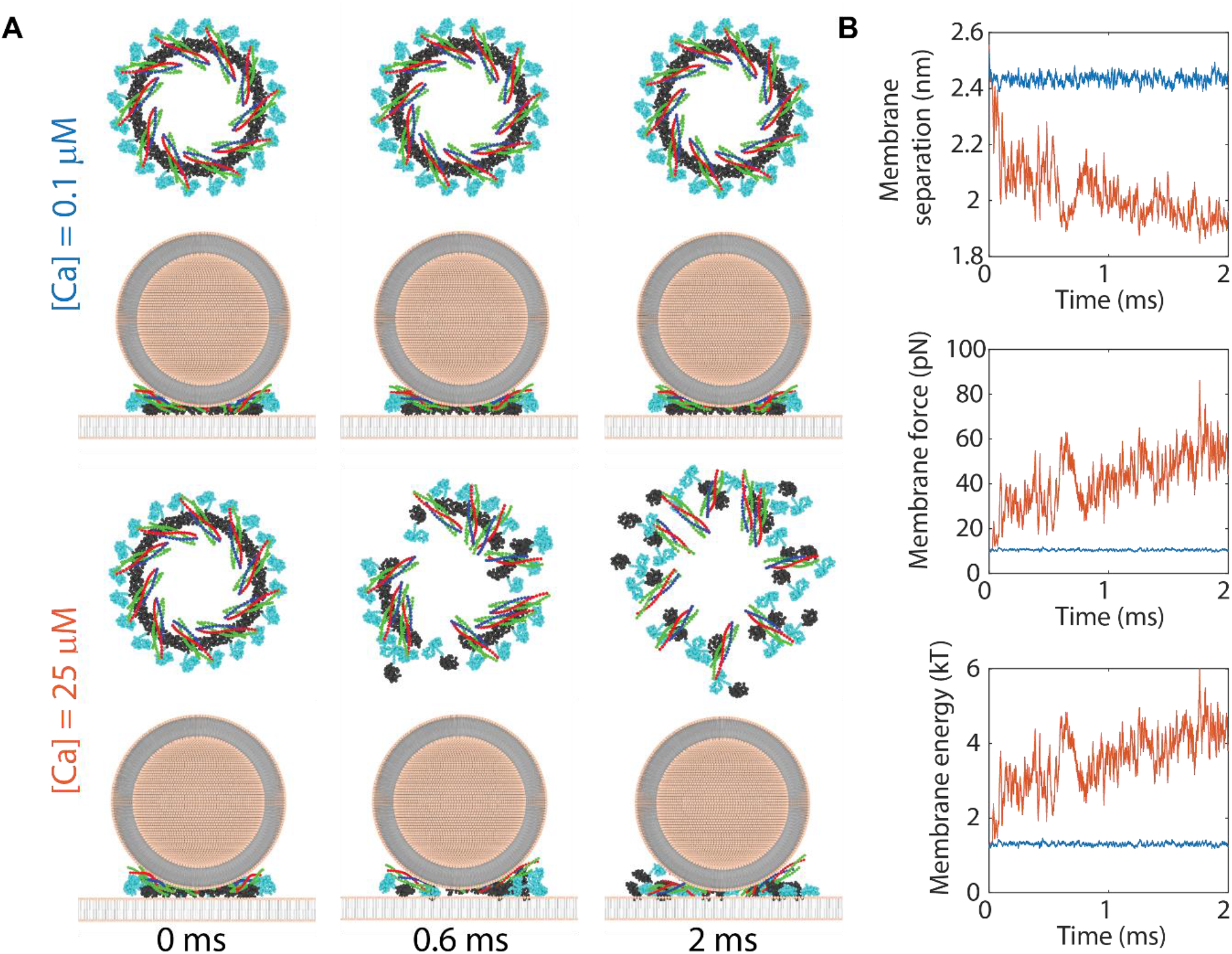
Syt rings form a Ca^2+^-sensitive fusion clamp. **(A)** Snapshots of simulations with basal (top two rows) and elevated (bottom two rows) presynaptic [Ca^2+^] values. With basal presynaptic [Ca^2+^], Syt rings were stable throughout the simulation, while with [Ca^2+^]=25 μM, they gradually disassembled, releasing the SNARE complexes. **(B)** Membrane separation (top), force (center), and energy (bottom) vs. time in typical simulations with basal (blue) and elevated (orange) presynaptic [Ca^2+^]. With basal [Ca^2+^], the Syt ring sterically separated the vesicle and PM, leading to low interaction forces and energies. With elevated [Ca^2+^], the Syt ring progressively disassembled, allowing the separation between the PM and vesicle to gradually decrease, increasing the interaction energy and force.

In simulations with basal [Ca^2+^], Syt rings formed a stable clamp, consistent with our previous study (35). Syt clamped fusion by imposing a separation between the vesicle and the PM of 2.4 ± 0.1 nm, large enough to keep the membrane energy below the threshold for fusion 1.2 ± 0.3 *kT*, Fig. 3A & B. The rings proved highly stable: in 100 simulations, each lasting 2 ms, the Syt ring remained intact, and no vesicles were fused. Syt rings also constrained the SNARE complexes, preventing them from assembling into a ring and fusing the vesicle and PM, as they would in the absence of the Syt clamp.

Ca^2+^ entry triggered rapid disassembly of the Syt ring, removing the steric block created by the ring. In simulations with 25 μM Ca^2+^, Syt-Syt bonds progressively dissociated as Ca^2+^ ions bound the C2B domains (16, 36), Fig. 3A. As the Syt ring disassembled, the separation between the vesicle and PM decreased, increasing the vesicle-PM interaction energy and the force pressing the membranes together, Fig. 3B. As observed in recent experiments and our previous simulations, Ca^2+^ binding triggered Syt-SNARE bond dissociation, because it is highly energetically favorable for Ca^2+^-bound C2B domains to insert their Ca^2+^-binding loops into the PM, a configuration that is sterically incompatible with binding to a fully zippered SNARE complex (35, 57).

Once released from the Syt ring, SNARE complexes drove rapid fusion. As in the SNARE-only simulations, SNAREs progressively self-assembled into a ring, Fig. 3A. Entropic forces then drove expansion of the SNARE ring, pulling the membranes together and increasing the membrane energy above the fusion threshold, Fig. 3B.

### Simulations starting from the unclamped state

The above results demonstrate that the Syt ring clamps fusion in a Ca^2+^-sensitive fashion. We next asked if the Ca^2+^ sensor Syt plays a somewhat less expected role in the membrane fusion function of the NT release machinery. Our SNARE-only simulations described above quantified the effects of SNAREs on the dynamics of membrane fusion, Fig. 2. However, experiments suggest that Syt may also increase the fusion rate at elevated intracellular Ca^2+^ concentrations in addition to its functional role as a clamp, helping to ensure release is tightly synchronized with incoming the Ca^2+^ stimulus (8, 13, 14, 58).

To isolate the possible fusion-related functions of Syt beyond its role as a Ca^2+^-sensitive clamp, we next performed simulations beginning in a completely unclamped state, varying the number of Syt monomers present at the fusion site. Vesicles were simulated for 1 ms with 100 μM Ca^2+^ prior to data logging to trigger complete unclamping. Syt enhanced membrane fusion by pulling the vesicle downward toward the PM via the linker domains. In simulations, the Ca^2+^-binding loops of Syt C2B domains rapidly penetrated into the PM, Fig. 4A. Since Syt is tethered to the vesicle via the linker domain, this binding creates a downward force on the vesicle, pulling it towards the PM. As a consequence, the force pressing the vesicle and PM together increased as a function of the number of Syt molecules at the fusion site, decreasing the fusion time, Fig. 4B & C. Similar to our measurements of the fusion time as a function of the number of SNARE complexes, we find that the fusion time decreased exponentially with additional Syt monomers,

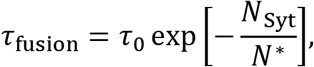

with best-fit value *N*\* = 4.89 ± 0.79 (determined using data with a range of numbers of SNARE complexes). This exponential scaling was valid independent of the number of SNARE complexes at the fusion site, Fig. 4C.

**Figure 4:**
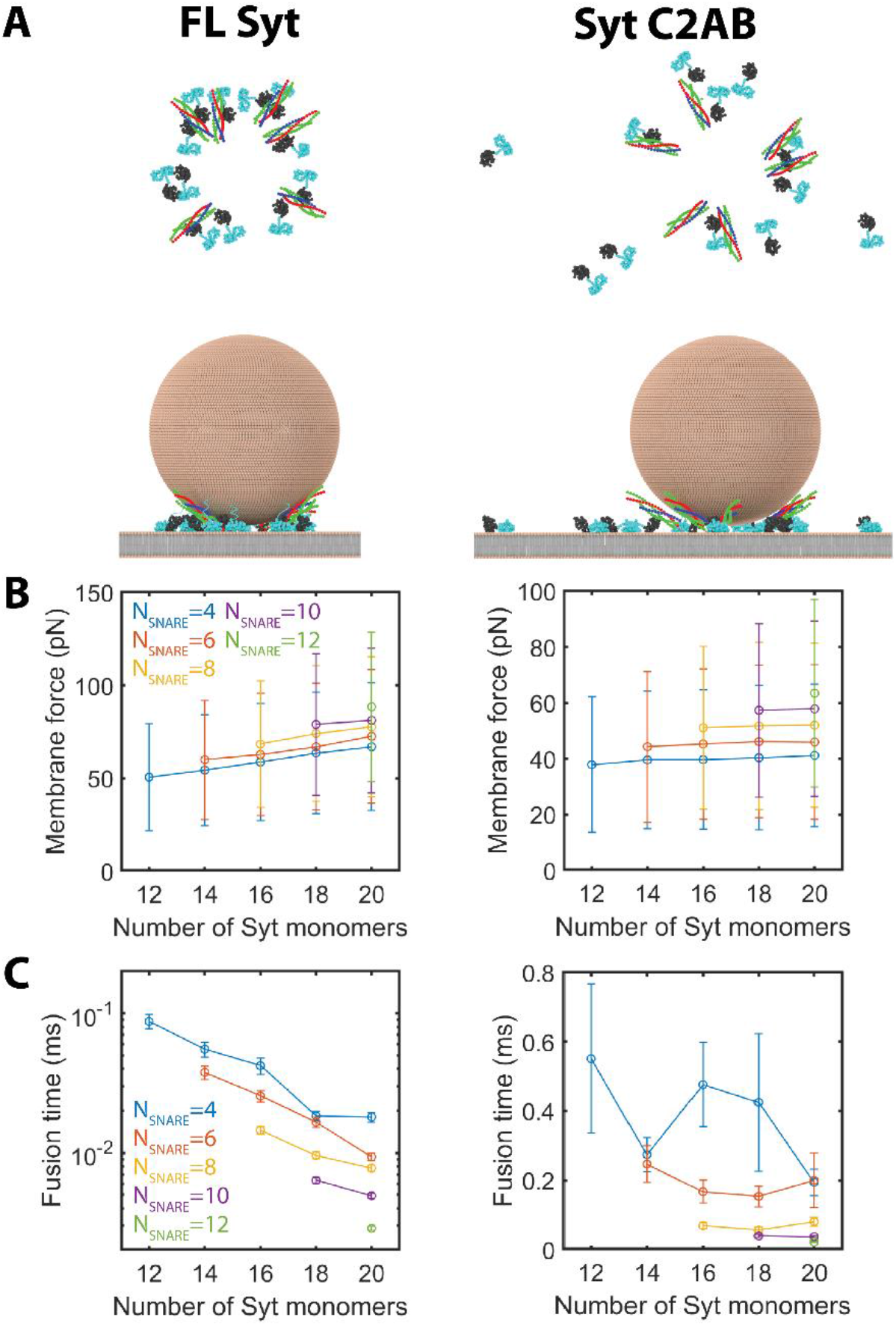
Syt activates fusion by tethering the vesicle to the PM. (**A**) Snapshots from simulations with 6 SNARE complexes and 12 Syt monomers with [Ca^2+^]=100 μM. Full length (FL) Syt molecules are tethered to the vesicle via the linker domain and are prevented from diffusing laterally along the PM (left), while C2AB Syt diffuses freely (right). Tension in the linker creates a downward force pulling the vesicle towards the PM. Membranes are depicted with explicit lipids, though simulations represented them as continuous surfaces. (**B**) Vesicle-PM force vs. the number of Syt monomers at the fusion site, with full length (left) and C2AB (right) Syt. FL Syt led to a pronounced increase in the force that grew with additional Syt monomers. Forces with C2AB Syt were lower, and did not increase with more Syt monomers. (**C**) Fusion time in the unclamped state vs. number of Syt molecules at the fusion site with full length (left) and C2AB (right) Syt. FL Syt increased the fusion rate relative to C2AB Syt. The fusion time decreased exponentially as a function of the number of FL Syt molecules. Additional C2AB Syt molecules did not systematically affect the fusion rate.

Enhancement of the fusion rate required the Syt C2AB domain to be tethered to the vesicle via the Syt LD. We performed simulations of C2AB Syt by disabling the force associated with the LD, so that the C2AB domains were detached from the vesicle. In these simulations, the membrane force was decreased relative to the force measured with full-length Syt, while the fusion time was increased, Fig. 4B & C. Furthermore, removal of the LD abolished the dependence of the membrane force and fusion time on the number of Syt molecules at the fusion site.

These results suggest that Syt-vesicle tethering contributes to synchronizing release with stimuli, because in the absence of this tethering, fusion times would be ~50 times larger, taking *N*_syt_ = 20. This is consistent with experiments in the Drosophila neuromuscular junction, which showed that the cytosolic fragment of Syt and Syt constructs tethered to the PM rather than to synaptic vesicles are not able to support synchronous NT release (59). Similarly, *in vitro* studies found that Syt-mediated activation of fusion requires membrane-anchored Syt to interact in *trans* with the target membrane by binding either anionic phospholipids or t-SNAREs (60).

### Increasing numbers of SNARE complexes at the fusion site increases the vesicular release rate

Experiments modulating the number of SNARE complexes present at the release site have shown that with additional SNAREs, release probability, EPSC amplitude, and the Ca^2+^ sensitivity of release are all increased (30–32). The mechanism underlying these effects is not known. While our SNARE-only simulations and previous results suggest the fusion rate is enhanced with more SNARE complexes, it is not clear how these effects would manifest in Ca^2+^-evoked NT release.

To address this question, we performed simulations with [Ca^2+^] fixed at a constant value in the range 5-40 μM, mimicking Ca^2+^ uncaging at the calyx of Held, a method in which [Ca^2+^] is elevated uniformly throughout an axon terminal (30, 50, 56, 61–63). We varied the number of assembled trans-SNARE complexes hosted by each vesicle, while keeping the number of Syt monomers fixed at *N*_Syt_ = 20. Again, Syt molecules were initially oligomerized into a ring. We predicted EPSCs evoked by Ca^2+^ uncaging at the calyx of Held by convolving the distribution of simulated vesicle release times (normalized by the number of vesicles in the readily-releasable pool at the calyx of Held) with a miniature excitatory post-synaptic current (mEPSC) signal (50).

Vesicles with more SNARE complexes released more rapidly, consistent with our observations above that increasing numbers of SNARE complexes accelerates fusion in the absence of Syt (28, 29), Fig. 2B & C. With presynaptic [Ca^2+^] elevated to 25 μM (*n* = 100 runs), s the number of SNARE complexes at the fusion site increased from 6 to 10, the mean release time decreased from 4.6 ± 1.9 ms to 2.5 ± 1.1 ms, Fig. S2.

The amplitude of excitatory post-synaptic current (EPSC) signals predicted by our simulations increased with additional SNARE complexes, independent of [Ca^2+^], Fig. 5A, suggesting that strengthened entropic forces caused by additional SNAREs enhance NT release. At high [Ca^2+^] (>~15 μM), the EPSC amplitude reached a limiting value, which was higher with more SNARE complexes.

**Figure 5:**
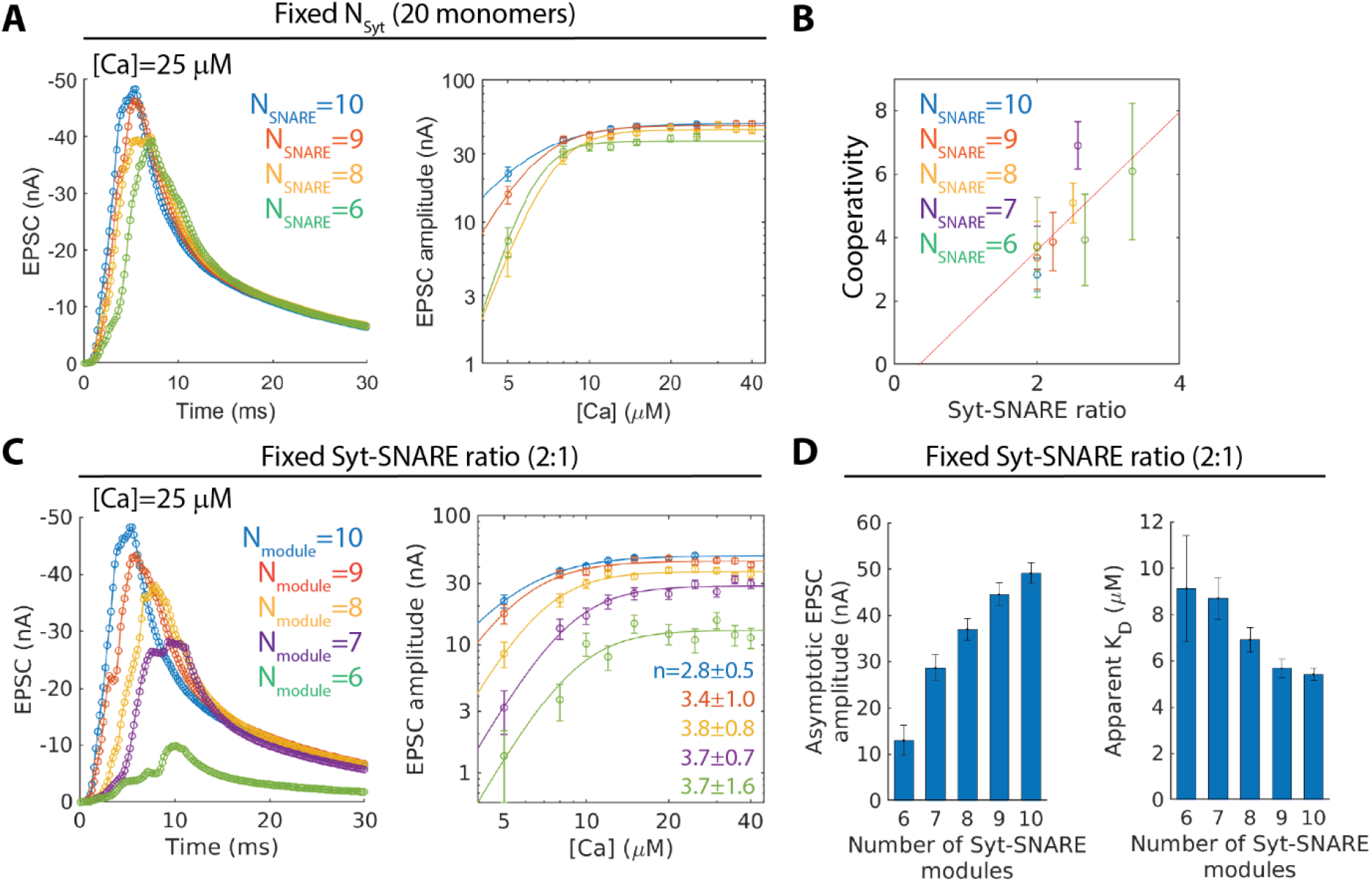
Composition of the vesicular release machinery determines cooperativity and sensitivity of NT release. **(A)** Ca^2+^ uncaging simulations varying the number of SNARE complexes with 20 Syt molecules at the fusion site. Left: simulation-predicted EPSCs with [Ca^2+^]=25 μM. Right: EPSC amplitude vs. [Ca^2+^]; curves represent best-fit Hill functions to the simulation data. Decreasing the number of SNARE complexes led to lower EPSC amplitudes, and steeper dose-response curves. **(B)** Best-fit Hill coefficients from dose-response curves measured varying the composition of release machinery. The Hill coefficient was well-fit by a linear relation with the stoichiometric ratio of Syt molecules to SNARE complexes (red line) (*R*^2^ = 0.56, *p* = 0.01, F-test). **(C)** Ca^2+^ uncaging simulations varying the number of Syt-SNARE modules, with 2 Syt molecules per SNARE complex. Left: simulation-predicted EPSCs with [Ca^2+^]= 25 μM. Right: EPSC amplitude as a function of presynaptic [Ca^2+^]; curves represent best-fit Hill functions to the simulation data. Varying the number of modules at the fusion site with fixed stoichiometry led to no change in the apparent cooperativity of the dose-response curve. **(D)** Maximal EPSC amplitude (left) and apparent dissociation constant *K*_D_ from best-fit Hill function fits to Ca^2+^ uncaging data. Additional Syt-SNARE modules increased the asymptotic EPSC amplitude dramatically and decreased the apparent *K*_D_.

### The cooperativity of NT release is determined by the stoichiometry of the release machinery

Decades of research have established a ~4^th^ power scaling of EPSC amplitude (or vesicle release rate) as a function of [Ca^2+^] as a hallmark of NT release at many diverse synapses (50, 64–66). Experiments showed that altering the number of SNARE complexes at the release site leaves the cooperativity of release was unchanged, but changes the Ca^2+^ sensitivity of NT release (30–32).

We measured the EPSC amplitude *A* as a function of [Ca^2+^] for a given number of SNARE complexes, and fit the result to a Hill function, 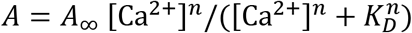, treating the apparent cooperativity *n*, the maximal EPSC amplitude *A*_∞_, and the apparent dissociation constant *K*_D_ as fitting parameters, Fig. 5A. We found that the apparent cooperativity of NT release varied depending on the number of SNARE complexes at the fusion site, with best-fit Hill coefficient values ranging from ~3-7, seemingly inconsistent with experiment (30–32).

However, we noted that the best-fit cooperativity *n* increased roughly linearly as a function of the stoichiometric ratio of Syt molecules to SNARE complexes, Fig. 5B. To test this, we performed additional Ca^2+^ uncaging simulations with 6-10 SNARE complexes and 12-20 Syt monomers at the release site, keeping *N*_SNARE_ ≤ *N*_Syt_/2. Resultant dose-response curves were also roughly consistent with a linear relation,

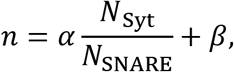

with best-fit values *α* = 22±0.7, *β* = −0.8±1.6.

Thus, based on the well-known apparent cooperativity of ~3-5, our simulations suggest that ~2-3 Syt monomers act to clamp each SNARE complex at the fusion site in neurons. Indeed, we found that fixing the Syt-SNARE ratio of 2:1 and varying the number of Syt-SNARE “modules” (each comprising two Syt molecules and one SNARE complex), the physiological cooperativity was reproduced roughly independent of the number of modules, Fig. 5C.

Acuna et al. (30) hypothesized that the observed changes in the dose-response function upon increasing the number of SNARE complexes at the fusion site were best explained by the assumption that the stoichiometry of the release machinery remained unchanged. Our results are consistent with this conclusion, and suggest that the stoichiometry of the release machinery was similarly unchanged in the experiments of Gerber et al. (31) and Arancillo et al. (31).

### The Ca^2+^ sensitivity of NT release and the asymptotic EPSC amplitude increase with additional Syt-SNARE modules

Experimental studies have shown that the Ca^2+^ sensitivity of NT release increases (defined as the inverse of the observed *K*_D_ of the dose-response curve) with the number of SNARE complexes at the fusion site (30–32).

In simulations, the apparent sensitivity of NT release to Ca^2+^ increased with the number of Syt-SNARE modules at the fusion site, reproducing the observations of refs. (30–32). With 10 modules, Hill function fits yielded an apparent dissociation constant of *K*_D_ = 5.4 ± 0.3 μM, while with 6 modules, the dissociation constant increased to *K*_D_ = 9.1 ± 2.3 μM, Fig. 5C. Thus, the Ca^2+^ sensitivity of NT release decreased by a factor of ~1.7.

Increasing the number of Syt-SNARE modules also increased the asymptotic EPSC amplitude at high [Ca^2+^], Fig. 5C & D. While increasing the number of SNARE complexes with a fixed number of Syt molecules led to a similar effect, Fig. 5A, the effect was much more dramatic at fixed stoichiometric ratio.

This effect can be readily understood. Fig. 2C & 4C show that additional SNARE complexes and Syt molecules each accelerate membrane fusion in the unclamped state. At high Ca^2+^ concentrations, the Ca^2+^-dependent unclamping step would be expected to occur rapidly, such that the later fusion step becomes rate-limiting (35), and therefore determines the time distribution of release events and the shape of the EPSC. Each additional Syt molecule and SNARE complex would accelerate the fusion step, increasing the EPSC amplitude. Increasing the number of Syt-SNARE modules adds Syt and SNAREs in parallel, dramatically enhancing NT release.

### Vesicle release probability increases with additional Syt-SNARE modules

We have shown that additional Syt-SNARE modules enhance release evoked by Ca^2+^ uncaging and increase the Ca^2+^ sensitivity of release. However, during physiological neurotransmitter release stimulated by an action potential (AP), the local [Ca^2+^] at the release site is only elevated transiently before rapid chelation by intracellular Ca^2+^ buffers such as parvalbumin and calretinin (67–69). How does the number of Syt-SNARE modules at the fusion site affect AP-evoked release? To address this question, we performed simulations of vesicles with varying numbers of Syt-SNARE modules using an AP-evoked [Ca^2+^] time course inferred from a kinetic model of NT release at the calyx of Held (56). We fixed the stoichiometric ratio of Syt monomers to SNARE complexes at 2:1.

Transiently elevated local [Ca^2+^] was insufficient to unclamp and drive release of most vesicles. During AP simulations, the majority of vesicles did not release, regardless of the number of assembled SNARE complexes at the fusion site, in agreement with experimental measurements showing a release probability per vesicle of ~0.1 at the Calyx of Held (70), Fig. 6A & 6B. Indeed, we found that during the AP-evoked [Ca^2+^] transient, most vesicles remained clamped: with 10 Syt-SNARE modules, the fraction of unclamped vesicles peaked at ~50% after ~0.25 ms, where we defined vesicle to be unclamped if at least 5 SNARE complexes are not bound to the Syt ring, Fig. 6C.

**Figure 6:**
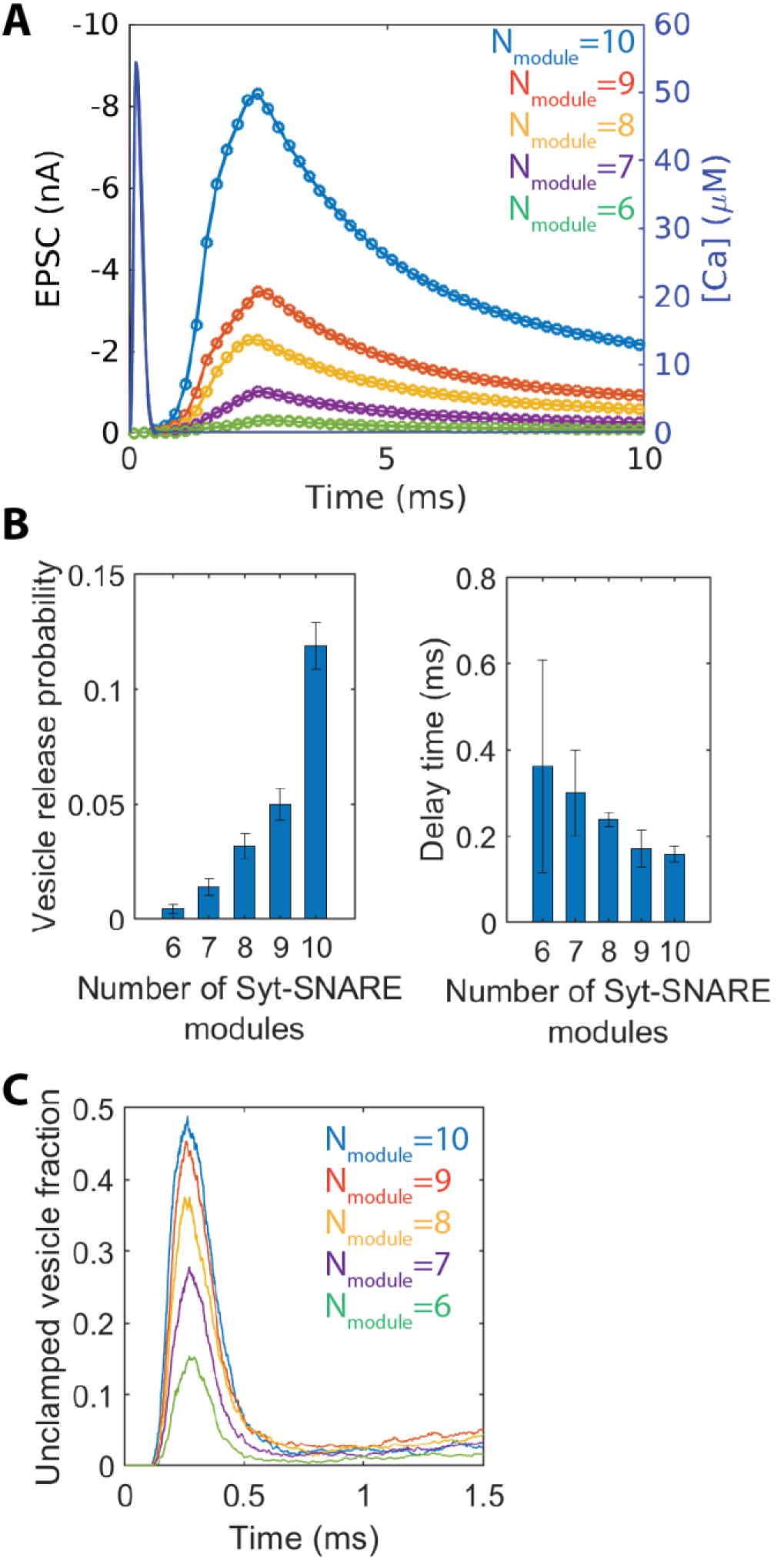
Addition of Syt-SNARE modules increases release probability. **(A)** Simulation-predicted EPSCs evoked by an action potential (AP) at the calyx of Held. With more Syt-SNARE modules, EPSC amplitude is dramatically increased in response to the same Ca^2+^ transient. **(B)** Vesicle release probability (left) and delay time (right) vs. number of Syt-SNARE modules. Vesicle release probability increased dramatically with additional modules, while the synaptic delay time decreased comparatively little. **(C)** Fraction of simulated vesicles that were unclamped (defined as having 5 or more SNARE complexes that were not bound to the Syt ring) as a function of time. Vesicles were unclamped transiently, with the unclamped fraction peaking at ~0.25 ms independent of the number of Syt-SNARE modules.

Increasing the number of Syt-SNARE modules at the fusion site increased the probability of release dramatically. With six modules, we found a release probability *p*_ves_ = 0.005 ± 0.002, while with 10 modules, the release probability increased to *p*_ves_ = 0.12 ± 0.01, Fig. 5B, reproducing the experimentally observed effects of modulating the number of SNARE complexes (30–32).

By what mechanism does increasing the number of Syt-SNARE modules at the fusion site increase release probability? The briefly elevated [Ca^2+^] caused transient unclamping of vesicles, Fig. 6C. Vesicle release can only occur during this brief unclamping window. Increasing the number of Syt-SNARE modules had little effect on the duration of the unclamping window. However, vesicle fusion is accelerated with more SNARE complexes or more Syt molecules; thus, vesicles with more modules are more likely to fuse during the transient unclamping episode.

### The synaptic delay time depends weakly on the number of SNARE complexes at the fusion site

To predict AP-evoked EPSCs, we again convolved the observed distribution of vesicle release times in our AP simulations with an experimentally measured mEPSC at the calyx of Held (50), repeating the procedure used in our Ca^2+^ uncaging simulations. We used these EPSCs to compare our results to commonly measured quantities in electrophysiological recordings.

Given that membrane fusion is faster with additional Syt-SNARE modules, it seems natural to suppose that increasing the number of modules will decrease the synaptic delay time, i.e. the time from Ca^2+^ entry to the start of the EPSC (in our analysis, the start of the EPSC was defined as the time when 2% of simulated vesicles had released). Surprisingly, we found that increasing the number of modules at the fusion site had only a weak effect on the delay time, Fig. 5C. With 10 Syt-SNARE modules, we found a delay time of 0.16 ± 0.02 ms, while the delay time with 6 modules was increased slightly to 0.36 ± 0.25 ms, Fig. 6B. Thus, lowering the number of Syt-SNARE modules from 10 to 6 increased the delay time by a factor of ~2, whereas the vesicle release probability decreased by a factor of ~30.

Why was the delay time weakly affected when the number of Syt-SNARE modules had such a dramatic effect on the fusion rate and release probability? The delay time measures the time for the first vesicles to fuse with the plasma membrane, which occurs shortly after the first vesicles become unclamped (35). Therefore, the duration of Ca^2+^-mediated unclamping is the primary determinant of the synaptic delay time, rather than time from unclamping to fusion. The unclamping time is largely unaffected by the number of Syt-SNARE modules hosted by the vesicle.

## Discussion

Here, we presented simulations of Ca^2+^-triggered NT release, including two central elements of the release machinery: the Ca^2+^ sensor Syt, which forms a Ca^2+^-sensitive clamp, and the core of the fusion machinery, the SNARE proteins. Our simulations suggest that SNARE complexes cooperate to fuse membranes via entropic forces (28, 29), Fig. 2. We found that Syt rings clamped release at basal [Ca^2+^] and removed the clamp in response to elevated [Ca^2+^], consistent with the common view that Syt functions as a Ca^2+^-sensitive clamp, Fig. 3. However, we also found that, after unclamping, Syt monomers contribute to SNARE-mediated fusion by enhancing the fusion rate via vesicle tethering forces, Fig. 4. Our simulations reproduced experimentally observed enhancements of Ca^2+^-evoked NT release when additional SNARE complexes are present at the release site, Fig. 5 & 6. Together, these results demonstrate that the Ca^2+^-sensing/clamping and membrane-fusing parts of the release machinery are functionally overlapping: the Ca^2+^ sensor contributes to accelerating membrane fusion, and the SNARE proteins contribute to the Ca^2+^-dependence of release.

### SNARE complexes and Syt monomers operate in modules with fixed stoichiometry

Our results suggest a picture in which SNARE complexes and Syt molecules do not operate independently to achieve distinct functions, but instead cooperate in modules for NT release, with 2-3 Syt molecules per SNARE complex. Experiments increasing or decreasing the number of SNARE complexes at the release site, respectively, showed increases or decreases in the Ca^2+^ sensitivity of NT release, with no change in the cooperativity (30–32); our simulations recapitulated this conservation of the cooperativity only if we assumed that the stoichiometric ratio of Syt molecules to SNARE complexes was conserved in these experiments. Increasing the number of Syt-SNARE modules enhanced release without altering the cooperativity, Fig. 5B & C, while increasing the number of SNARE complexes with a fixed number of Syt molecules decreased the Ca^2+^ cooperativity of NT release, Fig. 5A & B. The observed cooperativity of ~4 was reproduced with a Syt-SNARE ratio of 2:1.

Given that Syt rings comprise ~16 copies of Syt on average (17, 37), the stoichiometry predicted by our model suggests that ~5-8 Syt-SNARE modules per vesicle might regulate release in cells, consistent with cryo-EM reconstructions that have recently found six-fold symmetric protein structures under docked vesicles in PC12 cells and hippocampal neurons (71, 72). It is interesting to speculate that variations in the Syt-SNARE ratio could account for some of the variability of Ca^2+^ cooperativity of release at different synapses (73).

### The Ca^2+^ dose-response curve is an emergent property of the release machinery

Many pharmacological and genetic treatments that do not directly target the Ca2+-sensing and clamping components of the release machinery nevertheless alter the Ca^2+^ sensitivity of release. Lou et al. demonstrated an increase in Ca^2+^ sensitivity in cells treated with phorbol esters (61), which are thought to lower the energy barrier to fusion via a Munc13- and protein kinase C-dependent pathway (74, 75). Neurons treated with low concentrations of botulism neurotoxin A, botulism neurotoxin C1, or tetanus neurotoxin, also showed decreased Ca^2+^ sensitivity (33). Complexin knockout impaired both Ca^2+^- and sucrose-evoked release from mouse cortical neurons and induced a decrease in Ca^2+^ sensitivity (76). These effects demonstrate the apparent Ca^2+^ sensitivity of NT release is not solely controlled by the Ca^2+^-sensing and clamping components of the release machinery, but is also regulated by membrane-fusing components, showing that the membrane-fusing and Ca^2+^-sensing aspects of the release machinery cannot be conceptually separated. In this picture, the Ca^2+^ sensitivity is an emergent property of a complex system with many coupled components; enhancements in the membrane-fusing elements of this machinery increase the release rate at all Ca^2+^ concentrations, causing an apparent increase in the Ca^2+^ sensitivity.

The dose-response curves predicted by the present simulations showed that the asymptotic EPSC amplitude at high [Ca^2+^] increases with more Syt-SNARE modules, Fig. 5A & C. This effect was also observed in the experiments of Arancillo et al. (32): at high extracellular [Ca^2+^], EPSC amplitude was ~2-fold lower in cells with reduced Syntaxin expression compared to control cells, after correcting for the ~3-fold reduction of the readily-releasable pool (RRP). After correcting for changes in the RRP, this effect is not seen in the experiments of Acuna et al. (30). However, our simulations predict that increasing the number of Syt-SNARE modules from 8 to 9, for example, only increases the maximum EPSC amplitude ~20%, an effect that may be difficult to measure experimentally on top of a large change in the size of the RRP. Sakaba et al. (33) also found that the asymptotic release rate per vesicle decreased in cells treated with botulism neurotoxin A, though whether these effects can be attributed to a decrease in the number of Syt-SNARE modules mediating release is uncertain.

### Syt increases fusion rates by vesicle-PM tethering

Our simulations recapitulated the Ca^2+^-sensitive clamping function of Syt, but showed that in addition Syt enhances the fusion rate via an LD-dependent mechanism (4, 8, 13, 14, 18, 58, 77). This is an interesting intersection between the Ca^2+^-sensing and membrane-fusing components of the release machinery. In simulations with basal Ca^2+^ concentrations, Syt rings sterically separated the vesicle and PM, preventing the separation from reaching a sufficiently low value for fusion, Fig. 2. In contrast, in elevated Ca^2+^ concentrations, Syt monomers increased the fusion rate by pulling the vesicle towards the PM via the ~60-residue LD connecting the Syt TMD to the C2A and C2B domains. Deleting this linker domain abolished the effects of Syt on the fusion rate and the vesicle-PM interaction force, Fig. 4B & C, and Syt monomers were able to freely diffuse within the PM away from the vesicle, Fig. 4A.

Thus, the clamping components of the release machinery boost the membrane fusion rate by exerting force in a Ca^2+^-dependent manner. Following Ca^2+^ injection, membrane-bound Syt C2 domains that try to diffuse away from the vesicle create tension in the LD tethers that restrict them. This tension increases the force pressing the vesicle and plasma membranes together, which boosts the fusion rate. This model explains previous experimental findings that *trans* anchoring of the vesicle and target membranes is required for Syt to activate fusion (58, 59).

### Increased entropic forces explain the effects of additional SNARE complexes at the fusion site

It is commonly thought that the energy released by SNARE zippering is directly used to overcome the activation barrier to membrane fusion. Based on this reasoning, one might think that the number of SNARE complexes at the fusion site has little effect on the kinetics of release, and instead simply determines whether fusion can be achieved. A more detailed mathematical model based on similar reasoning argued that the fusion rate depends non-monotonically on the number of SNARE complexes at the fusion site, with 4-6 SNARE complexes maximizing the fusion, motivating the hypothesis that this number may be selected in neurons (51). It is not clear how these models can be reconciled with electrophysiological studies showing that release is progressively enhanced with additional SNARE complexes and impaired with fewer SNARE complexes (30–32), suggesting that release is accelerated by increasing the number of SNARE complexes.

When SNARE complexes, vesicles, and plasma membranes are explicitly simulated, an entirely different picture emerges, in which membrane fusion is driven by force that originates from SNARE-SNARE and SNARE-membrane entropic forces (28, 29). The conclusion that SNARE-mediated fusion is driven by entropic forces is consistent with the gradual, monotonic enhancement of NT release with additional SNARE complexes demonstrated by many experimental studies. Fusion can then be achieved by any number of SNARE complexes, but is exponentially faster with more SNAREs. Additional SNAREs accelerate fusion by enhancing entropic forces that push the SNAREs outward, pulling the vesicle towards the PM, as described in previous studies (28, 29, 35). A natural outcome is that additional SNAREs enhance release rates.

### The membrane fusion machinery controls release probability due to transient unclamping

While many mechanisms are known by which the Ca^2+^-sensing machinery regulate release probability, the effects on release probability of the membrane-fusing aspects of the machinery are less well-characterized (78). Our simulations showed enhanced release probability with more Syt-SNARE modules, Fig. 6B, consistent with experiment (30–32). This follows form the fact that transient elevation of the presynaptic Ca^2+^ concentration transiently unclamps vesicles, Fig. 6C. In order for a vesicle to release, it must fuse with the PM during the window of time in which it is unclamped. Increasing the number of Syt-SNARE modules increases the release rate during this unclamping window, leading to a larger number of released vesicles. A similar effect may also drive experimentally observed changes in release probability upon treatments expected to change the energetic barrier to fusion, such as application of phorbol esters, or mutations altering the surface charge density of SNARE complexes (79, 80): lowering the energy barrier to fusion increases the fusion rate, and hence the release probability (35).

## Acknowledgements

This work was supported by National Institute of General Medical Sciences of the National Institutes of Health under award number R01GM117046. The content is solely the responsibility of the authors and does not necessarily represent the official views of the National Institutes of Health. We acknowledge computing resources from Columbia University’s Shared Research Computing Facility project.

