## Supporting Information for "SNAREs and Synaptotagmin cooperatively determine the Ca^2+^ sensitivity of neurotransmitter release in fixed stoichiometry modules"


Zachary A. McDargh[1], Ben O'Shaughnessy[1]*

[1]Department of Chemical Engineering, Columbia University, New York City, NY-10027, USA

### Analytical model of SNARE zippering kinetics

Our model is based on two assumptions: (1) SNARE complexes zipper independently and (2) individual SNARE complexes zipper with a constant probability per unit time $k$. Given an initial condition in which all SNARE complexes are un-zippered, we determine the mean time at which a collection of $N_{\mathrm{SNARE}}$ SNARE complexes becomes zippered.

Based on assumption (2), the probability that a given SNARE has not zippered at time $t$ is an exponential, given by

$$s(t) = \exp(-k\ t).$$

From assumption (1), the probability that all $N_{\mathrm{SNARE}}$ SNARE complexes *have* zippered at time $t$ is therefore given by

$$c(t) = (1 - \exp(-k\ t))^{N_{\mathrm{SNARE}}}.$$

This can also be interpreted as the cumulative distribution function for the time at which zippering is complete (i.e., when the last SNARE to zipper does so). We can therefore obtain the probability distribution function for the zippering time by differentiating with respect to $t$,

$$p(t) = k\ N_{\mathrm{SNARE}} \exp(-k\ t)\ (1 - \exp(-k\ t))^{N_{\mathrm{SNARE}}-1},$$

which immediately gives the expectation value of the zippering time,

$$\langle t \rangle = \int_0^\infty t\ p(t) \mathrm{d}t = k\ N_{\mathrm{SNARE}} \int_0^\infty t\ \ \exp(-k\ t)\ (1 - \exp(-k\ t))^{N_{\mathrm{SNARE}}-1} \mathrm{d}t = \frac{H_{N_{\mathrm{SNARE}}}}{k},$$

where $H_n$ indicates the $n$th harmonic number. The harmonic numbers are given by

$$H_n = \sum_{m=1}^{n} \frac{1}{m}.$$

For large $n$, the harmonic numbers are well approximated by

$$H_n \approx \gamma + \log n,$$

where $\gamma \approx 0.577$ is the Euler-Mascheroni constant. We use this approximation to fit this simplistic model to our simulation data. Fig. S1 shows a comparison of our simulation data, along with a best-fit function of the form

$$\langle t \rangle = \frac{\gamma + \log N_{\mathrm{SNARE}}}{k},$$

treating $k$ as a fitting parameter. The model captures the slow increase of the zippering time as a function of $N_{\mathrm{SNARE}}$ very accurately.

**Table S1: Model Parameters**

| **Parameter** | **Value** |
|---|---|
| $\delta t$, Time Step of MD simulation | 0.1 ns |
| $\eta$, Viscosity of water | $8.9 \times 10^{-3}$ dyn · s/cm$^2$ |
| $T$, Temperature | 298 K |
| $\lambda_D$, Debye length[a] | 0.8 nm |
| $A$, Hamaker constant[b] | $8 \times 10^{-21}$ J |
| $l_p$, Persistence length of worm-like chain[c] | 0.6 nm |
| Contour length per residue of uncomplexed regions[c] | 0.365 nm |
| $\lambda_{hyd}$, Decay length for Hydration force[d] | 0.194 nm |
| $P_0$, Pressure prefactor for Hydration Force[d] | 44,273 $k_B T$/nm |
| Syt-SNARE capture distance | 0.7 nm |
| Syt-Syt capture distance | 1.5 nm |
| Syt-membrane capture distance | 0.8 nm |
| Syt-SNARE dissociation time | 25 µs |
| Syt-Syt dissociation time | 1010 µs |
| C2A-membrane dissociation time[e] | 7.4 ms |
| C2B-membrane dissociation time[e] | 2.0 s |
| C2AB-membrane dissociation time[e] | 60 s |
| Synaptotagmin Ca binding rate, $k_{on}$[f] | $2\times10^8$ M$^{-1}$s$^{-1}$ |
| Ca binding affinity of sites 1, 2, and 3 on C2A[g] | 120 µM, 465 µM, 1700 µM |
| Ca binding affinity of sites 1 and 2 on C2B[g] | 20 µM |

**Table S1.** (a) Calculated from physiological salt concentration of 0.15 M (1). (b) Typical for lipid bilayers (1). (c) Typical for unstructured polypeptides (2, 3). (d) Determined from composition-weighted averages (Table S4) of values measured for pure lipid species (4). (e) Off-rates measured in reference (5). (f) Tuned to reproduce synaptic delay times at Calyx of Held; similar to on rates in the literature (6) (7-9). (g) C2A values taken from ref. (10); C2B values tuned to reproduce probability of release at calyx of Held (see ref. (11)). The value used in our simulations is in the range of values affinities measured in the literature (10, 12, 13).

**Table S2: Bead charges in CG model of SNARE complex**

| | VAMP | Syntaxin | SN1 | SN2 |
|---|---|---|---|---|
| 1 | - | - | 0 | - |
| 2 | - | 0 | 1 | - |
| 3 | - | 0 | -2 | - |
| 4 | - | 2 | 2 | - |
| 5 | - | -1 | -1 | 0 |
| 6 | - | 0 | -2 | -2 |
| 7 | 2 | -1 | 0 | -2 |
| 8 | 0 | 0 | 1 | 0 |
| 9 | -1 | 0 | -2 | 0 |
| 10 | -2 | -1 | 0 | 1 |
| 11 | 1 | -1 | 1 | -1 |
| 12 | 0 | 0 | 0 | -1 |
| 13 | 0 | -1 | -2 | -1 |
| 14 | 0 | -2 | -2 | 0 |
| 15 | -2 | 0 | -1 | 0 |
| 16 | -1 | -1 | 0 | 0 |
| 17 | 0 | -1 | -1 | 2 |
| 18 | 0 | 0 | 0 | -2 |
| 19 | -1 | -1 | 0 | 1 |
| 20 | 1 | 2 | -1 | -1 |

**Table S2.** Charges in units of the electronic charge $e$. Bead 20 represents layer +8 of the SNAREpin, with bead numbers decreasing towards the N-terminus. Each bead comprises four residues, with charge determined by the total charge of the constituent residues at pH 7, calculated using the all atom crystal structure PDB ID 3HD7 (14) and the CHARMM forcefield (15).

**Table S4: Membrane composition**

| | Vesicle | Plasma membrane |
|---|---|---|
| PE | 23% | 40% |
| PC | 36% | 40% |
| PS | 12% | 18% |
| PIP2 | 0 | 2% |

**Table S4.** Vesicle composition taken from (16). Plasma membrane composition taken from (17).

### SI Figures

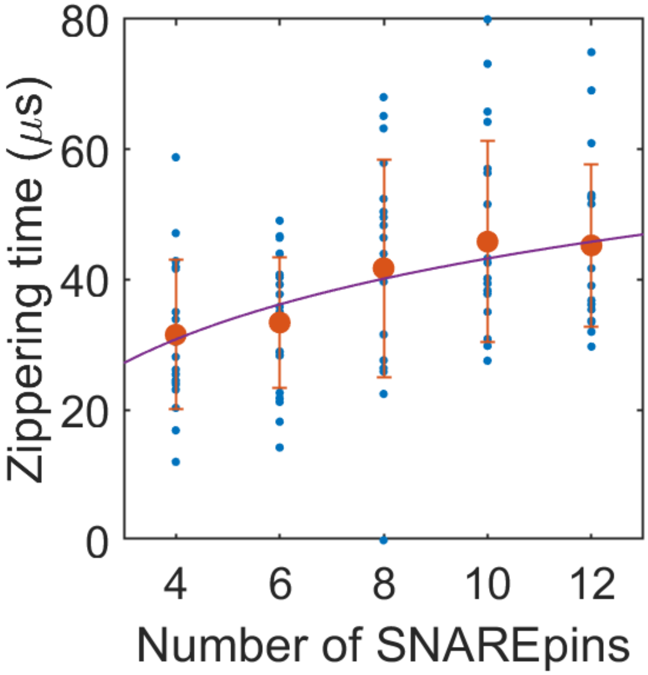


**Figure S1: SNARE zippering time increases weakly with additional SNARE complexes at the fusion site**

Time until all $N_{\text{SNARE}}$ SNARE complexes are fused as a function of $N_{\text{SNARE}}$ in simulations lacking Syt. Blue points represent individual simulations, orange points represent the mean zippering time with a given number of SNARE complexes (error bars are SD). A purple curve shows the predicted zippering times from an analytical model that assumes all SNARE complexes zipper independently; the model includes one fitting parameter, the average time for one SNARE complex to zipper. Least-squares fit of the simulation data to the analytical prediction gave a zippering time for a single SNARE of $14.9 \pm 1.0$ μs.

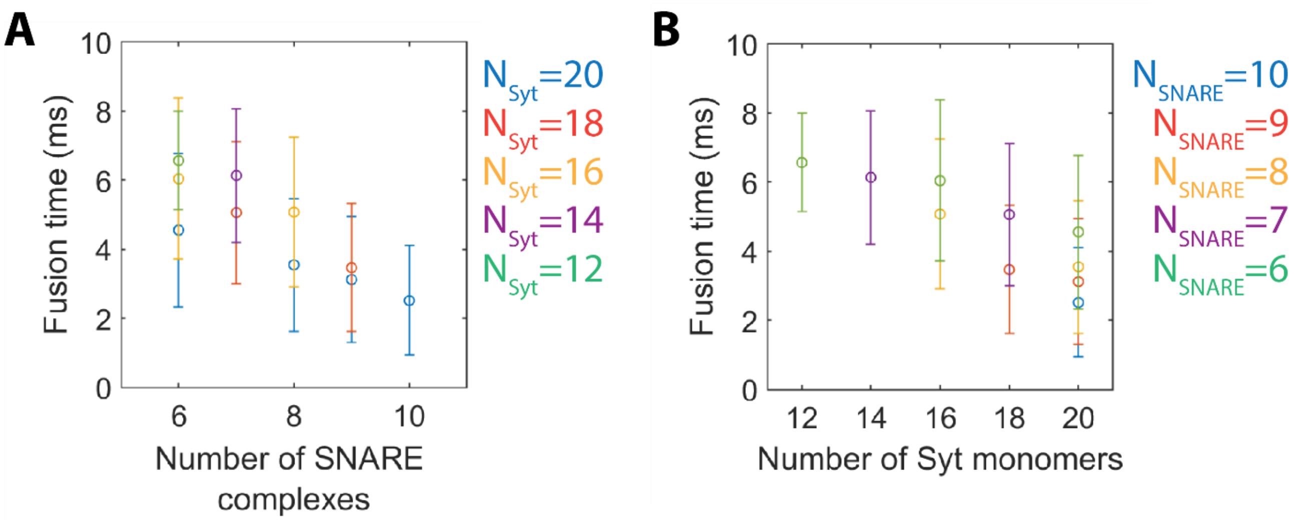


**Figure S2: Fusion is faster with more SNARE complexes and more Syt monomers**

**A)** Average release time in simulations of vesicles with the indicated number of SNARE complexes and Syt monomers at the fusion site, with the Syt molecules initially assembled into a ring. These simulations were performed with [Ca] = 25 μM.

**B)** The same data contained in panel (A), replotted to clearly show the dependence of the fusion time on the number of Syt monomers at the fusion site.
